# Nearly unbiased estimator of adult population size based on within-cohort half-sibling pairs incorporating flexible reproductive variation

**DOI:** 10.1101/422659

**Authors:** Tetsuya Akita

## Abstract

Close-kin mark-recapture (CKMR) is a kinship-based method for estimating adult abundance. However, the application of CKMR is limited to using a kinship relationship that is not affected by family-correlated survivorship, which leads to a biased estimation. We developed a nearly unbiased estimator of the number of mothers in a population, which is based on the known maternal half-sibling relationship found within the same cohort. Our method allowed for variance of the averaged offspring number per mother (between-age variation) and for variance of the offspring number among mothers with the same reproductive potential (within-age variation). Estimators of its variance and coefficient variation were also provided. The performance of the estimators was quantitatively evaluated by running an individual-based model. Our results provide guidance for (i) a sample size to archive the required accuracy and precision when the order of mother size is available and (ii) a degree of uncertainty regarding the estimated mother size when information about the mother size is not available. Taken together, these findings offer an opportunity to shed light on the usefulness of analysing within-cohort half-sibling pairs and will greatly widen the scope of the CKMR method.

## Introduction

Estimating population size is a fundamental component of wildlife management and stock assessment. Molecular markers can provide information about both absolute population size and effective population size. The former is a direct measurement of biomass in a population and is thus of interest especially in stock assessment, while the latter reflects the degree of genetic diversity (Wright 1931). To estimate absolute population size, mark-recapture (MR) studies using genetic information via tissue sample collection have several advantages over traditional methods, including noninvasive sampling (Luikart *et al.* 2010).

Close-kin mark-recapture (CKMR) is a recently developed method that uses information about relatedness in a sample, made possible by recent advances in genetic methods for kinship determination (Bravington *et al.* 2016a,b; Skaug 2017; Hillary *et al.* 2018), although similar ideas have been proposed in the beginning of the 21st century (Nielsen *et al.* 2001; Pearse *et al.* 2001; Skaug 2001). The rationale is that the presence of a kinship pair in the sample is analogous to the recapture of a marked individual in MR. Kinship pairs in the sample are less likely to be found in larger populations; thus, the number of kinship pairs may reflect adult abundance. The first large-scale application of CKMR was the estimation of the absolute abundance of southern bluefin tuna based on the detection of 45 parent–offspring pairs (POPs) in 13,000 samples (Bravington *et al.* 2016a). CKMR relaxes the methodological constraints of MR and allows for the following: a single sampling occasion of an individual (or tissues) is sufficient; and the sampling of offspring provides information about their parent number. These features are suitable for applications in fishery management, such as by-catch and large-scale commercial fisheries.

At the current stage of CKMR, POPs and half-sibling pairs (HSPs) have been successfully used for estimating adult population size. HSP-based CKMR, which involves many more DNA markers than POP-based CKMR, has recently become applicable due to cost reduction. Although the use of HSPs in combination with POPs is more informative in estimating adult population size (CCSBT 2017), using only HSPs has the practical advantage of not requiring adult samples (Hillary *et al.* 2018). However, when HSPs are sampled from the same cohorts (referred to as “within-cohort HSPs”), a larger variation in offspring number per parent leads to a higher probability of HSP in the pair, resulting in a significantly biased estimation of adult population size (Bravington *et al.* 2016b, Akita 2018). This may be problematic when applying HSP-based CKMR to species such as marine species that have a type-III survivorship curve (i.e., big litters and a high early-life-stage mortality) showing overdispersed reproduction (Hedgecock and Pudovkin 2011; Eldon *et al.* 2016). There may be two ways to avoid this. The first is to exclude within-cohort HSPs from the estimation and only include pairs that are composed of different cohorts (referred as to “cross-cohort HSPs”), as is currently done in the stock assessment of southern bluefin tuna (CCSBT 2017) and in the estimation of the population size of white sharks (Hillary *et al.* 2018). This procedure can remove within-year fluctuations in individual reproductive success, such as family-correlated survival (Ottmann *et al.* 2016). However, it imposes additional constraints on the applicability of the HSP-based CKMR method. For example, cross-cohort HSPs are not found in semelparous species; the probability that two pairs (born in year *t*_1_ and *t*_2_) share a half-sibling relationship includes information about the adult population numbers at *t*_1_ and *t*_2_, resulting in inconvenient outputs for the annual index of the adult population. The second way to avoid a large bias in estimating adult population size is to design a theory that includes the effect of overdispersed reproduction. This method relaxes the constrains noted above and may greatly widen the scope of HSP-based CKMR, although the degree of overdispersion is generally unknown.

Here, we propose a new method for estimating the number of mothers in a population. This approach is based on the number of maternal half-sibling (MHS) pairs found within the same cohort–the one containing within-cohort HSPs–and on modelling that explicitly incorporates overdispersed reproduction, assuming that kinships are genetically detected without any errors. Our model divides the effect of overdispersion into two types of variations: (i) age- or size-specific differences in mean fecundity (referred to as “between-age variation”), and (ii) unequal contributions by mothers of the same age or size to the number of offspring at sampling (referred to as “within-age variation”). It should be noted that, when applying within-cohort HSPs to the estimation method of a population size, the latter type of variation is a key component, while the former type of variation affects the methods using both within- and between-cohort HSPs. First, we formulate the distribution of offspring number under the two types of variations. Second, we analytically derive the probability that two randomly chosen individuals found in the same cohort share an MHS relationship. Third, we find a nearly unbiased estimator of mother size and its relative estimators. Finally, we investigate the performance of the estimators by running an individual-based model. Our modelling framework may be applied to diverse animal species; however, the description of the model focuses on fish species, which are currently the best candidate target of this method.

## Theory

All symbols used in this paper are summarized in Table 1.

**Table 1:** List of mathematical symbols in the main text

|  |  |
| --- | --- |
| $n$ | Sample number of offspring |
| $n_{\text{pair}}$ | Number of pairs in a sample ($= {}_n\text{C}_2$) |
| $N$ | Number of mothers in the population when sampled offspring are born |
| $N_e$ | Effective number of mothers in the population |
| $\phi$ | Overdispersion parameter under negative binomial reproduction |
| $\lambda_i$ | Expected number of surviving offspring of the mother $i$ at sampling |
| $k_i$ | Number of surviving offspring born to the mother $i$ |
| $H$ | Number of maternal–half-sibling pairs found in samples |
| $\pi$ | Probability that a randomly chosen pair (two offspring) share a maternal half-sibling relationship |
| $c$ | Combined effect of deviation from the Poisson<br>($= (1 + \phi^{-1})\mathbb{E}[\lambda^2]/\mathbb{E}[\lambda]^2$) |
| $\hat{N}_0$ | Moment estimator of $N$ |
| $\hat{N}_1$ | Nearly unbiased estimator of $N$ |
| $\hat{v}$ | Estimator of $\mathbb{V}[\hat{N}_1]$ |
| $\hat{c}v$ | Estimator of $\mathbb{CV}[\hat{N}_1]$ |
| $b_{\text{mean}}$ | Bias of $\hat{N}_1$ |
| $b_{\text{var}}$ | Bias of $\hat{v}$ |

### Hypothetical population and sampling scheme

Suppose that there is a hypothetical population consisting of *N* mothers and that there is no population subdivision. In this paper, the mature female is referred to as a mother even if she does not produce offspring. We introduce the concept of reproductive potential, which is determined by several factors, including the mother’s age, weight, or residence time on the spawning grounds. The reproductive potential is allowed to be variable among mothers. In this work, this variation is referred to as between-age variation for the sake of better understanding; however, other factors besides age can be incorporated into the variation. In addition, the variation in reproduction among mothers with the same reproductive potential, referred to here as within-age variation, is also incorporated into the modelling, resulting in a large variation in the fertility of the mothers. Figure 1 provides an illustration of kinship relationships between mothers and their offspring. The mother with a higher reproductive potential is more likely to produce a larger number of offspring (i.e., between-age variation), but even with an identical reproductive potential, there is a large variation in offspring number (i.e., within-age variation).

**Figure 1.**
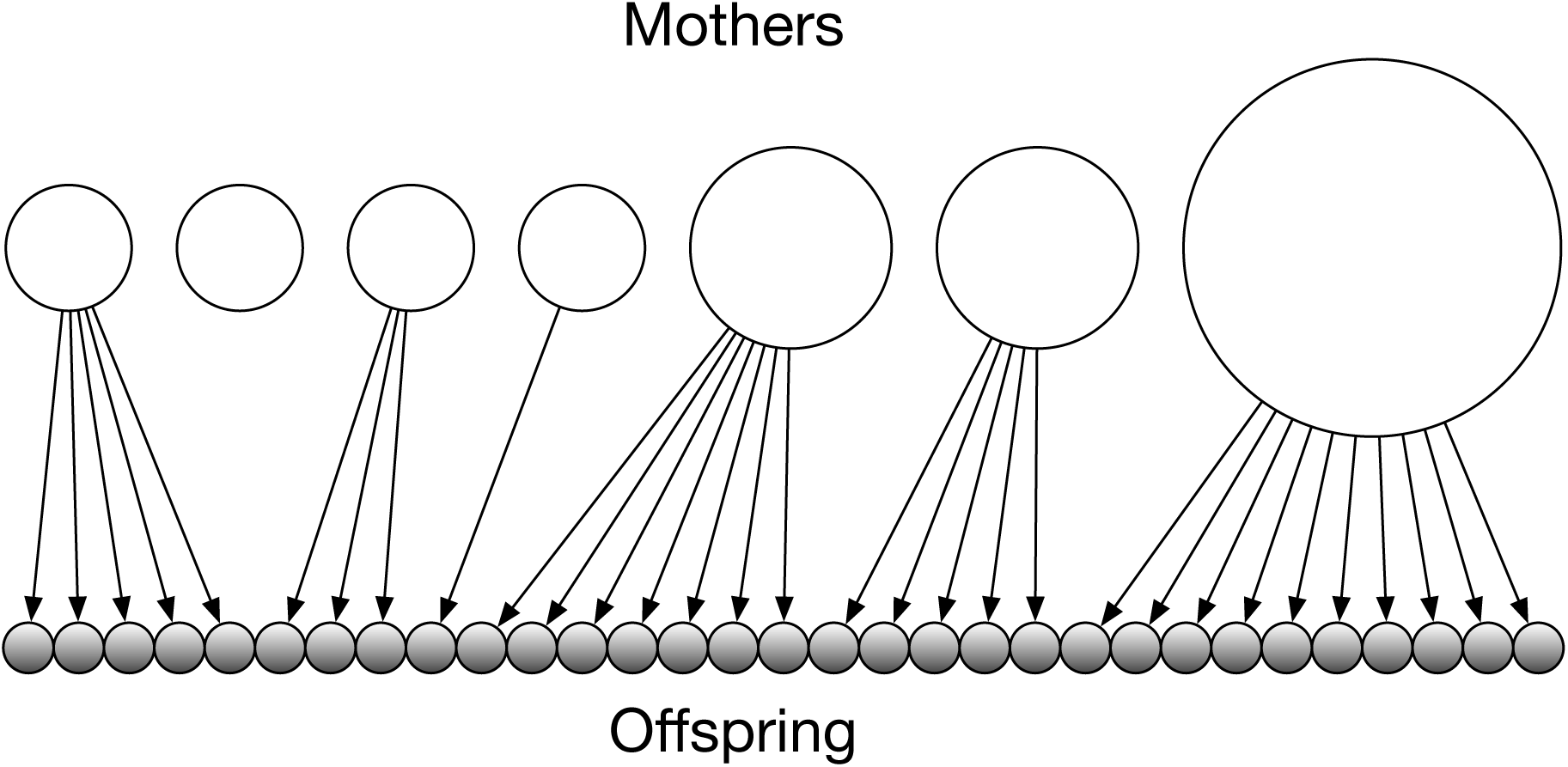
Example of relationships between mothers and their offspring number with *N* = 7. An open and closed circle indicate mothers and their offspring, respectively. The area of an open circle indicates the strength of reproductive potential for each mother. Arrows show mother–offspring relationships.

For the detection of MHS pairs, *n* offspring within the same cohort (i.e., same age) are simultaneously and randomly sampled in the population. For mathematical tractability, we assume that there is only one spawning ground and that the mothers will stay there during the whole spawning season. The number of offspring in the population varies with sample timing: the earlier the sampling, the larger the number of surviving offspring per mother, and thus, the larger the number of HSPs likely to be found in the sample. At each sample timing, we introduce the expected number of surviving offspring of mother *i* (*i* = 1, 2, *…, N*), *λ*_*i*_. This parameter summarizes the reproductive characteristics of mother *i* and can be considered the reproductive potential, as mentioned above. It should be noted that the magnitude of the parameter includes information about the survival rate of the offspring, the number of days after the eggs have hatched, and the egg number, which implies that the parameter reflects the sample timing.

Let *k*_*i*_ be the number of surviving offspring of mother *i* at sampling. Given the expected number of offspring *λ*_*i*_, *k*_*i*_ is assumed to follow a negative binomial distribution by a conventional parametrisation,

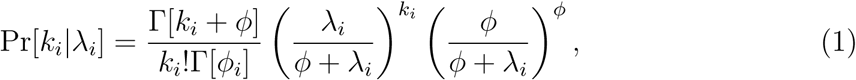

where *ϕ* (*>* 0) is the overdispersion parameter describing the degree of within-age variation (Akita 2018). At present, *ϕ* is assumed to be constant among mothers, whereas the expected number of surviving offspring (*λ*_*i*_) is variable among mothers. The mean and variance of this distribution are *λ*_*i*_ and 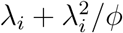, respectively. In the limit of infinite *ϕ*, this distribution becomes a Poisson distribution:

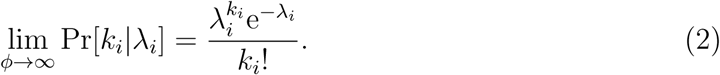

We assume that *λ*_*i*_ is independent and identically distributed with a density function *f* (*λ*), which produces between-age variation. The shape of the density function is often complex but may be described by information such as the mother’s weight composition. The specific form of *f* (*λ*) is given in the supporting information (see supplementary Appendix A) and used for checking the theory developed in the paper. As explained later, the theory does not require this specific form but only requires the ratio of the second moment to the squared first moment (i.e., 𝔼[*λ*^2^]*/*𝔼[*λ*]^2^).

### MHS probability among randomly chosen individuals

Here, we derive the probability that two offspring share an MHS relationship with an arbitrary mother. Given the realized number of offspring *k*_1_, *k*_2_, *…, k*_*N*_, the probability that two randomly chosen offspring are born to mother *i* is 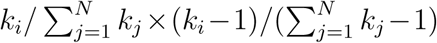. Hence, the conditional probability that two offspring share an MHS relationship with an arbitrary mother is

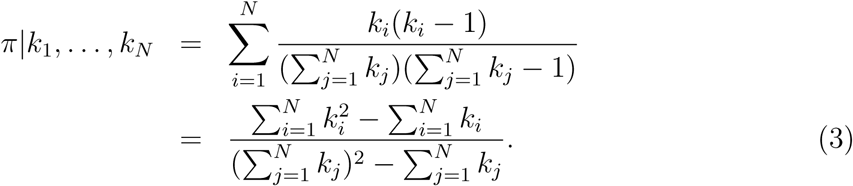

It should be noted that *k*_*i*_ is a random variable followed by a negative binomial distribution (Equation 1), where the parameter of the distribution, *λ*_*i*_, is also a random variable followed by an arbitral function *f* (*λ*). By taking the expectation over the distribution of offspring number, the following conditional probability is given by

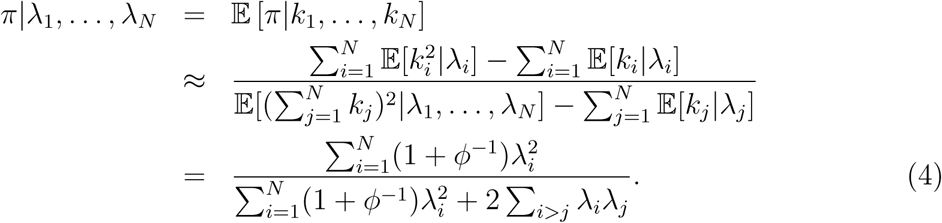

Going from the first line to the second line, we use the approximation that 𝔼[*g*_1_(*k*)*/g*_2_(*k*)] *≉* 𝔼[*g*_1_(*k*)]*/*𝔼[*g*_2_(*k*)]. The expectations average over *k* or *k*^2^, conditional on *λ*. By taking the expectation over *λ* and applying a similar approximation, the unconditional probability is given by

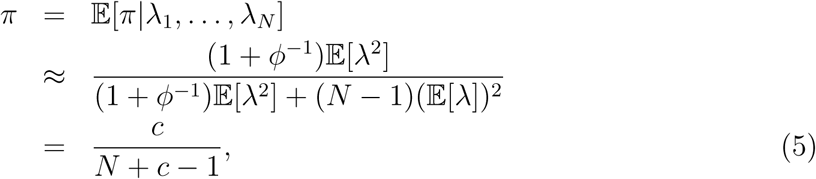

Where

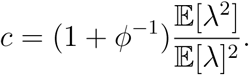

In computing the expectation, we drop the subscript (“*i*” or “*j*”), since *λ* is independent and identically distributed. Equation 5 explicitly indicates the two variations (i.e., between-age variation and within-age variation) that determine the degree of deviation from the Poisson. When *λ* is constant among mothers, *π* becomes (1 + *ϕ*^-1^)*/*(*N* + *ϕ*^-1^), which appears in Equation 7 of Akita (2018). In addition, as *ϕ → ∞*, (1+*ϕ*^-1^)*/*(*N* +*ϕ*^-1^) converges to 1*/N*, which corresponds to the Poisson variance of *k*_*i*_ for all mothers in a population. The effect of the two factors causing a deviation from the Poisson can be combined as a parameter *c* (*≥* 1); hereafter, “overdispersion” is referred to as the state of the offspring number distribution resulting from this combined effect.

When *N* is given, *π* increases with an increase in *c*, suggesting that a randomly chosen pair is more likely to share an MHS relationship under stronger overdispersion. Figures 2a and b show the theoretical curve and the simulation results of *π* as a function of *ϕ* and 𝔼[*λ*^2^]*/*𝔼[*λ*]^2^, respectively. For the investigated function *f* (*λ*), the approximation in Equation 5 is confirmed to work well.

**Figure 2.**
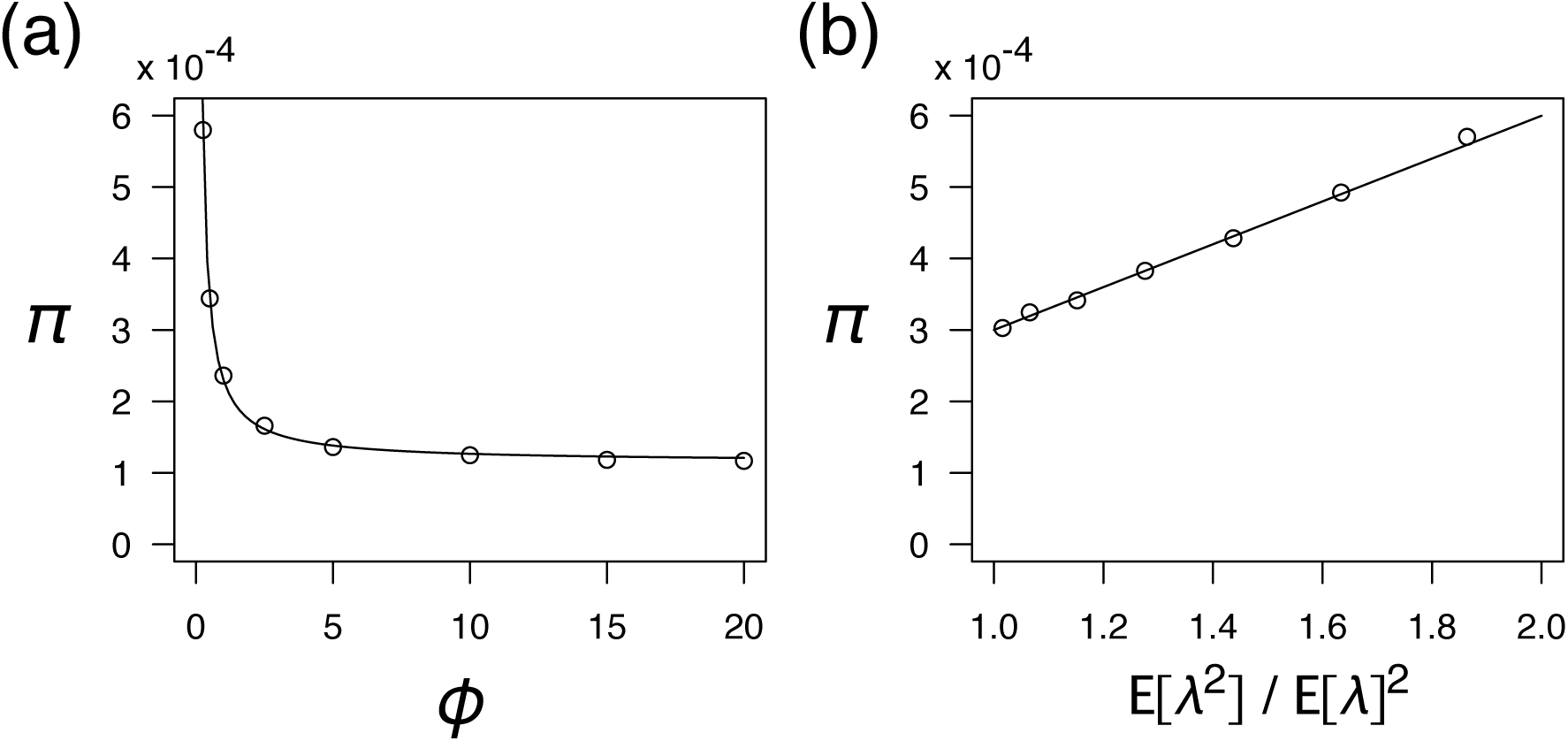
Accuracy of the approximation for *π* as a function of (a) *ϕ* and (b) E[*λ*^2^]*/*E[*λ*]^2^. The thin curve indicates the approximated values (Equation 5) and points indicate the simulated value from 10,000,000 replications. 𝔼[*λ*^2^]*/*𝔼[*λ*]^2^ in (b) are calculated from the distribution of *λ* (see supplementary Appendix A) with *b* = 0.3, 0.6, 0.9, 1.2, 1.5, 1.8 and 2.1. (a) *b* = 0.9. (b) *ϕ* = 0.5. Other parameters: *N* = 10,000. The definition of parameter *b* is described in supplementary Appendix A.

As noted in the previous subsection, the expected number of surviving offspring per mother (*λ*) is affected by the sample timing. However, the MHS probability (*π*, in Equation 5) is independent of sample timing, which is not involved in family-correlated survivorship. It is easily confirmed by using the relationship 𝔼[(*βλ*)^2^]*/*𝔼[*βλ*]^2^ = 𝔼[*λ*^2^]*/*E[*λ*]^2^, where *β* is a constant term involved in a family-uncorrelated sample timing.

### Statistical properties of the number of MHS pairs

In this subsection, given the unconditional probability that two offspring share an MHS relationship (Equation 5), we consider the distribution of the number of MHS pairs and its statistical properties. Let *H* be the number of MHS pairs found in an offspring sample of size *n*. First, we derive the approximate distribution of *H* for a situation where overdispersion does not exist (i.e., *c* = 1). Second, we evaluate the applicability of the derived distribution of *H* for the non-overdispersed situation to the overdispersed situation (i.e., *c >* 1).

If overdispersion does not exist (i.e., *c* = 1), drawing an MHS pair from a randomly chosen pair in a sample is considered a Bernoulli trial. Thus, *H* follows a hypergeometric distribution, which is a function of the sample size of the offspring, the total number of offspring in the population, and the total number of MHS pairs in the population. However, in the setting of this work, the latter two components are random variables, thus creating a complex situation for deriving the exact formulation (Akita 2018). Therefore, assuming that the total number of MHS pairs in the population is much larger than the number of pairs in a sampleΣ 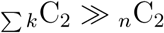 the distribution is approximated by a binomial form:

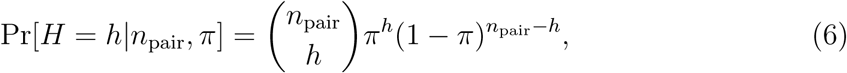

where *n*_pair_ is the number of pairs in a sample (= _*n*_C_2_). For practical purposes, the condition 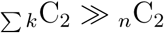 may be acceptable. The theoretical expectation of *H* is

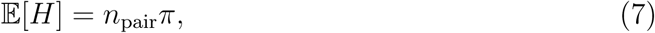

and the variance is

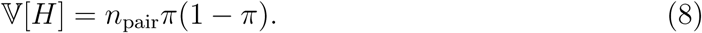

Figures 3a and c illustrate the accuracy of the theoretical prediction for the expectation and the variance of *H* under the Poisson variance as a function of *n*, respectively. For the investigated parameter, the prediction is confirmed to be quite accurate.

**Figure 3.**
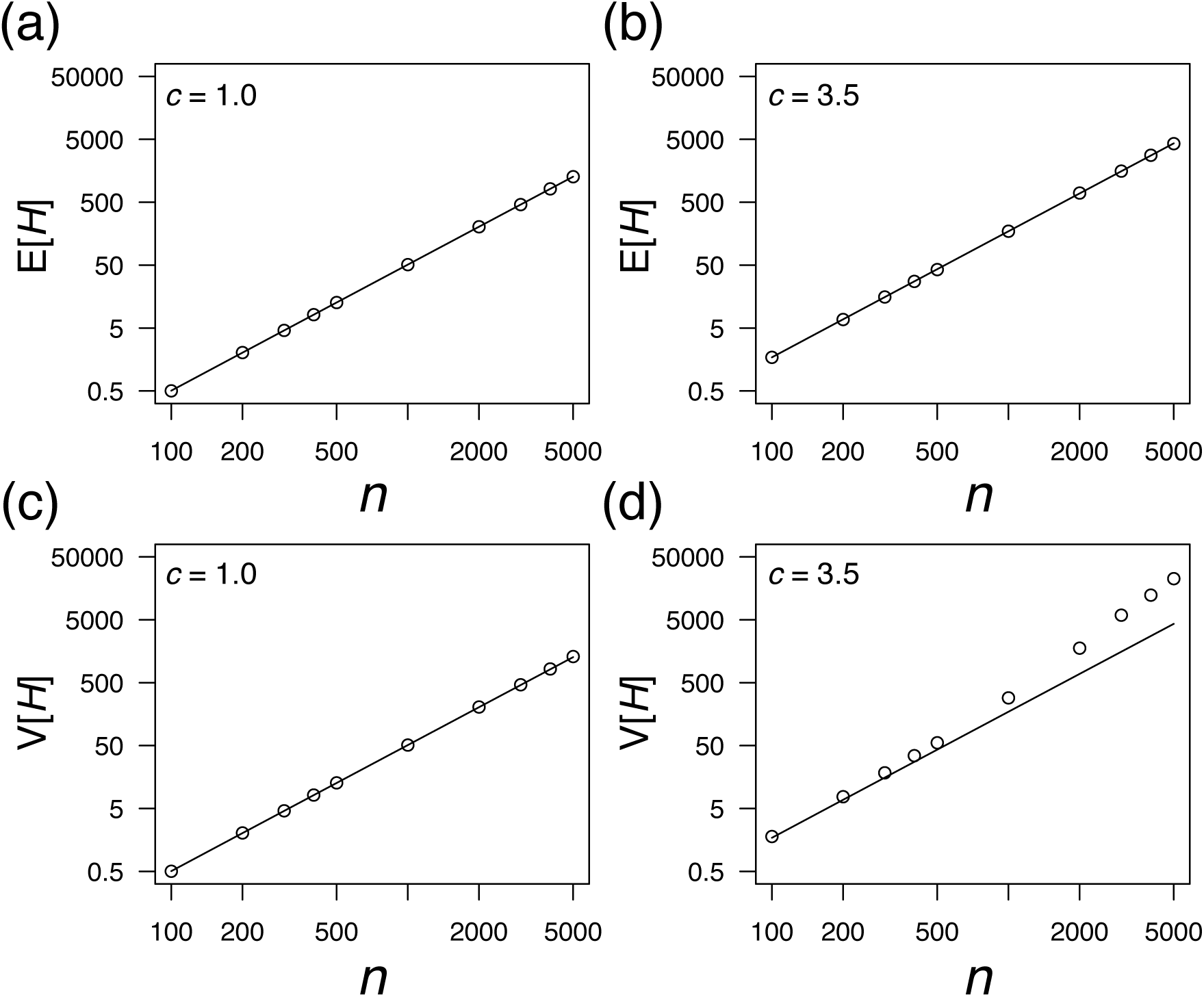
Accuracy of the theoretical prediction for 𝔼[*H*] (a, b) and V[*H*] (c, d) as a function of *n*. Thin lines indicate theoretical values (Equations 7 and 8) and points are obtained from simulated data (1,000,000 replications). The value of *c* is described on the panel. *N* = 10,000. Both *x*-axis and *y*-axis are log-scale.

If overdispersion exists (i.e., *c >* 1), drawing an MHS pair is no longer a Bernoulli trial. For example, an individual that is born to a relatively “successful” mother has a greater probability of an MHS relationship with other individuals. Therefore, it is expected that a hypergeometric/binomial form is not appropriate for the distribution of *H*. As shown in Fig. 3d, the binomial variance (Equation 8) is downwardly biased from the observed variance of *H* when *n* increases. The theoretical evaluation is relatively complex and is left for future research. However, for the investigated parameter set, the expected value is well approximated by Equation 7 (Fig. 3b), assuming independent comparisons. The rationale may be that the MHS probability in a pair, *π* (Equation 5), includes the effect of overdispersion. Next, on the basis of an accurate approximation of E[*H*] in the overdispersed situation, we provide the estimator of *N* from the observed number of MHS pairs in a sample.

### Estimator of the number of mothers from the observed number of MHS pairs

By removing *π* in Equations 5 and 7, *N* can be written as a function of *c, n*_pair_, and E[*H*]. The observed number of MHS pairs in a sample is defined by *H*_obs_ and E[*H*] is replaced by *H*_obs_, generating the moment estimator of *N*:

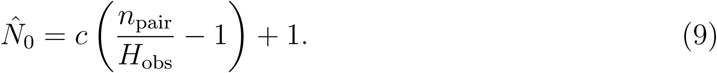

In this paper, the “hat” indicates the estimator of the variable. Assuming that *H* follows a binomial distribution, the estimator corresponds to the maximum likelihood estimator of *N* (see supplementary Appendix B). There are two problems with using this estimator. First, the value of 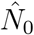 becomes inflated when there are no observations of MHS pairs ina sample (i.e., *H*_obs_ = 0). This leads to an inconvenient situation where an individual-based model (IBM) frequently generating zero MHS pairs is not available for statistical evaluation. Second, even if an MHS pair is found in a sample, 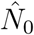 will be upwardly biased (see supplementary Appendix B). Therefore, an improved estimator is needed for the purpose of appropriate evaluation and higher accuracy for a wide parameter range.

In a similar spirit to Chapman (1951), we derived an alternative estimator of *N* (see supplementary Appendix C):

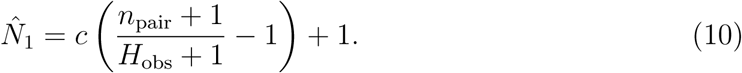

The bias of 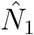 is defined by *b*_mean_, given by

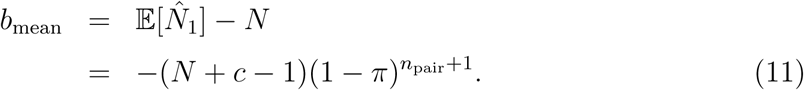

It should be noted that 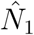 will be downwardly biased, although the bias may be ignored for a wider range of parameters than 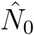 (see details in the Results section).

We also found the estimator of 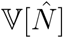, given by

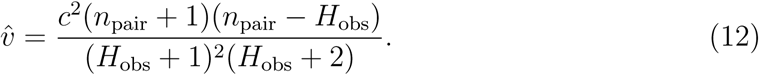

The derivation process is similar to Seber (1970) (see Appendix D for details). The bias of 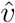 is defined by *b*_var_, given by

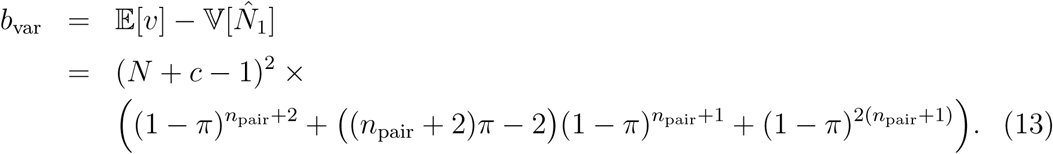

Finally, we consider the estimator of the coefficient of variation of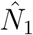. A method similar to deriving 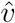 (i.e., searching the formula such that its expectation approximates 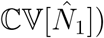 is very complex for the estimator; instead, using Equations 10 and 12, we defined the estimator by

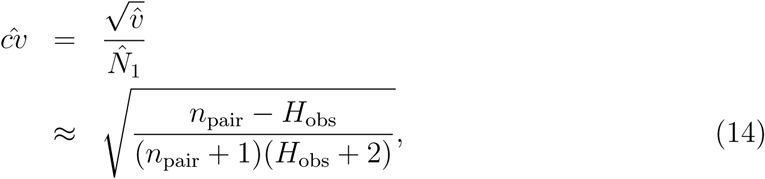

where 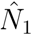is assumed to be *c*(*n*_pair_ + 1)*/*(*H*_obs_ + 1). It should be noted that, if this assumption is allowed, 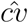is independent of *c*. Roughly speaking, 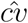 is approximated by 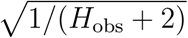 since *n*_pair_ *» H*_obs_, which is similar to an approximate lower bound on the coefficient of variation, as appears in Bravington *et al.* (2016b).

### Individual-based model (IBM)

To evaluate the performance of the estimators 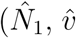 and 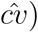, we developed an IBM that tracks kinship relationships. The population structure was assumed to be the same as in the development of the estimators. The population consisted of mothers and their offspring and was assumed to follow Poisson or negative binomial distributions. The expected number of surviving offspring of a mother followed the density distribution *f* (*λ*) (see supplementary Appendix A). It should be noted that the overdispersion parameter (*c*) was calculated from *ϕ* and *f* (*λ*). Each offspring retained the ID of its mother, enabling us to trace an MHS relationship.

Given a parameter set (*N, n, ϕ*, and parameters that determine *f* (*λ*)), we simulated a population history in which *N* mothers generate offspring, and this process was repeated 100 times. For each history, the sampling process was repeated 1000 times, acquiring 100,000 data points that we used to construct the distribution of 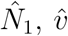 and 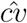 for each parameter set.

## Results

We evaluated the performance of the estimators 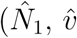 and 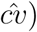 in a situation where the combined effect of deviation from the Poisson, *c*, was known. The parameter values were changed for *N* (10^4^ and 10^5^) and *c* (1, 1.4, and 3.5). We mainly addressed how many samples (*n*) would be required to achieve adequate accuracy and precision under a given parameter set (*N* and *c*).

First, we evaluated the accuracy of 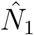 based on its bias (*b*_mean_). Given *N*, the absolute value of the bias is shown in Figs. 4c and 4d as a contour plot. For comparison, the results of 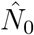 are also presented (Figs. 4a and 4b). Clearly, the absolute value of the bias is smaller in 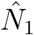 than in 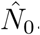. Hereafter, we use 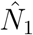 as the estimator of mother size.

**Figure 4.**
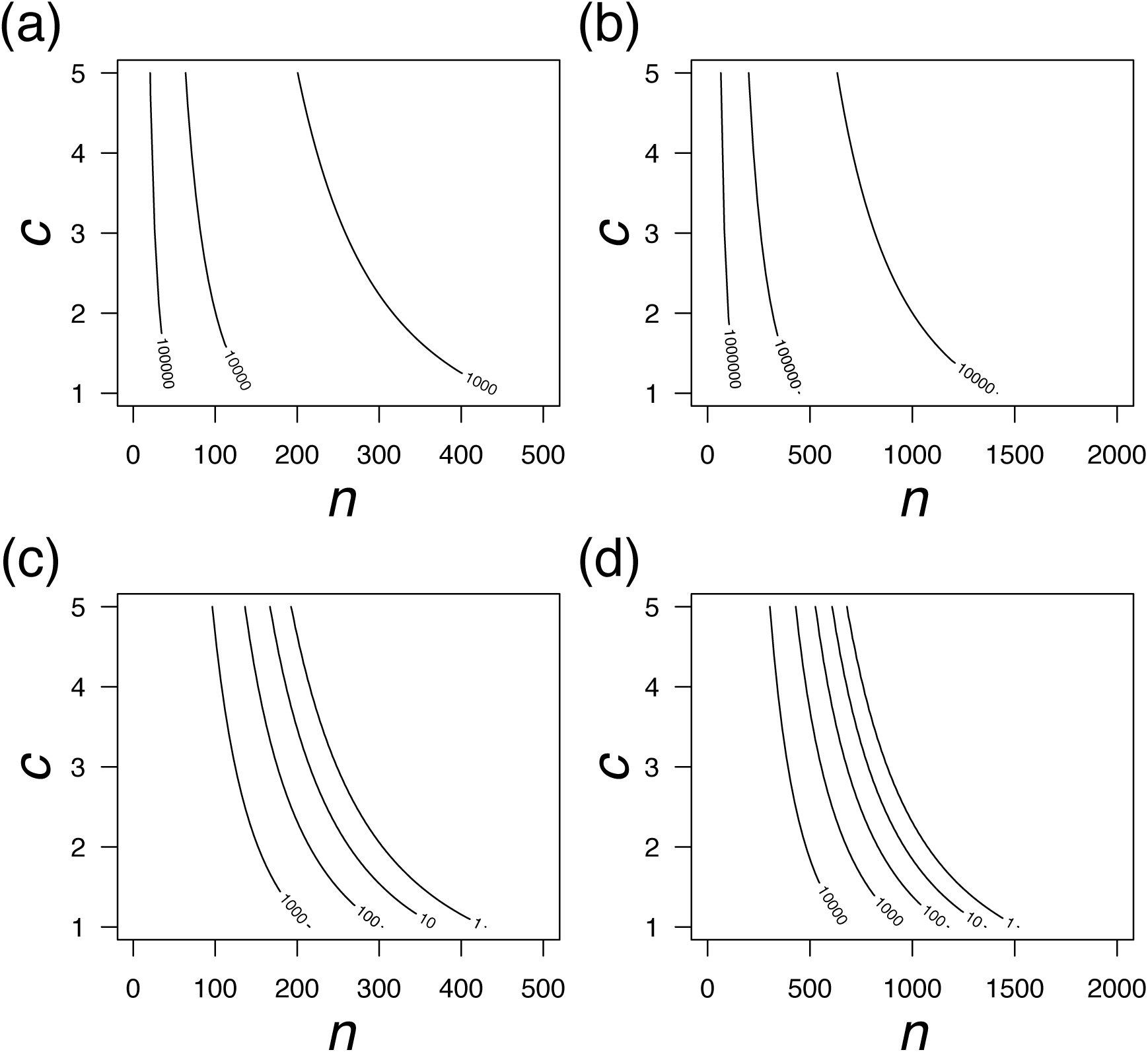
Contour plot of the bias of 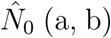 and 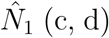 (c, d) as a function of *n* and *c*. The absolute value of the bias is shown on the panel. (a, c) *N* = 10,000. (b, d) *N* = 100,000.

In the situation of Poisson variance (i.e., *c* = 1), the requisite sample size (*n*) with a small bias, less than 1%, is approximately 300 for *N* = 10,000 (*|b*_mean_*| <* 100, Fig. 4c) and 1000 for *N* = 100,000 (*|b*_mean_*| <* 1000, Fig. 4d). As *c* increases, the range of *n* satisfying the requirement (e.g., *|b*_mean_*|/N <* 0.01) expands, since a larger *c* makes the term (1 *-π*)^*n*pair+1^ in *b*_mean_ vanish more quickly. Regardless of the value of *c, |b*_mean_*|* exponentially increases with a decrease in *n*, leading to a situation where the estimation of *N* is strongly downwardly biased when *n* is relatively small. When *n* is fixed, a larger value of *c* leads to a smaller value of *|b*_mean_*|*. The result of the IBM supported the above consideration. Figure 5 illustrates the averaged value of 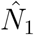 (denoted by open circles) with a 95% confidence interval (CI), which is obtained from the IBM. As expected, the averaged value of 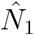 downwardly deviates from *N* given a relatively small sample size (*n*) satisfying *|b*_mean_*| »* 1. As *n* increases, the averaged value of 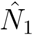 approaches a true *N* (denoted by a dotted line in Fig. 5).

**Figure 5.**
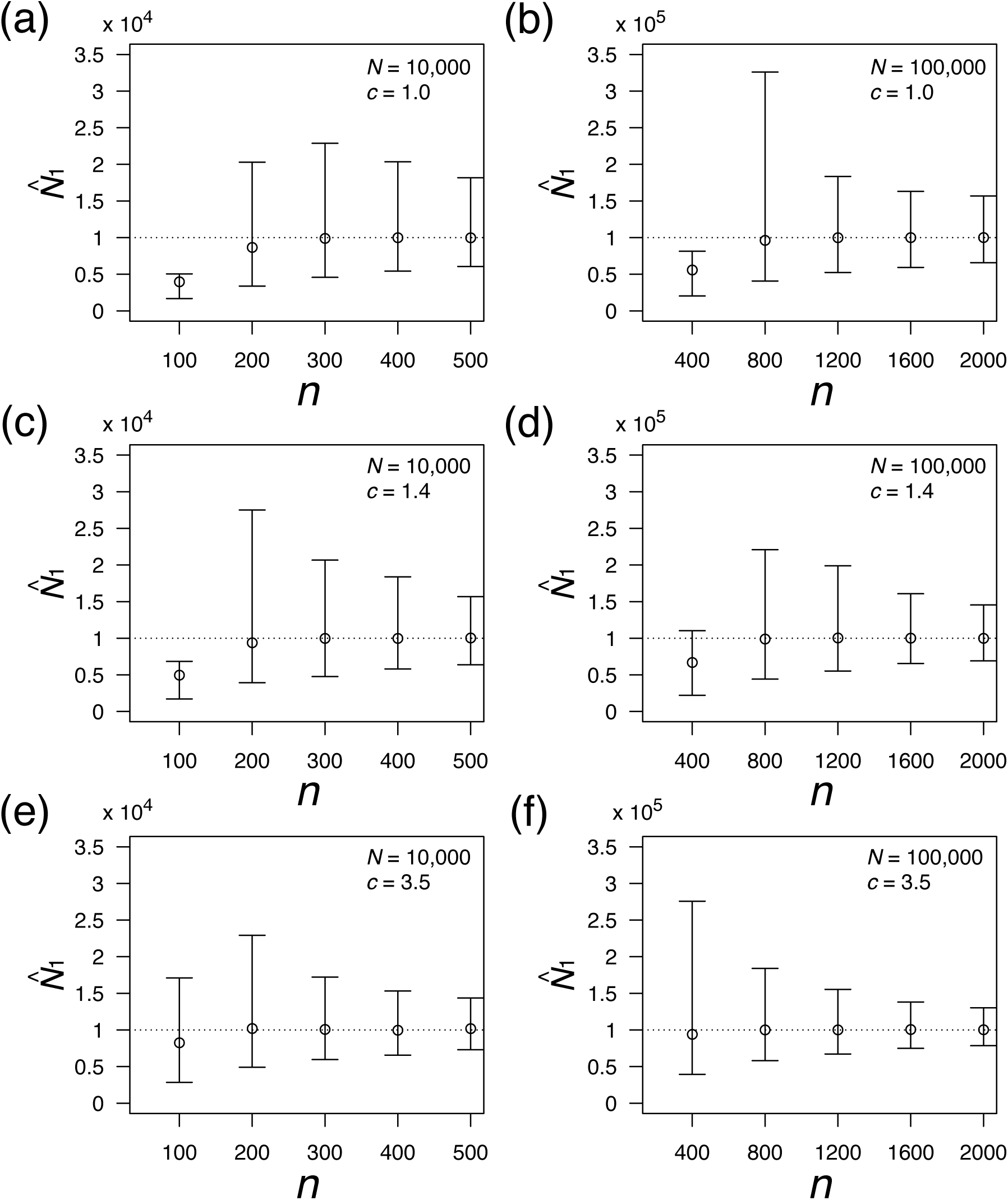
Accuracy and precision of 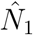1 as a function of *n*. Open circles represent means with 95% confidence intervals. A dotted line indicates the true value of *N*. The value of parameters (*N* and *c*) is described on the panel.

Second, we evaluated the precision of 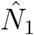. As shown in Fig. 5, the precision of 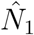 with a change in *n* behaves in a complex manner. For the investigated parameter set, while the lower limit of the CI monotonically increases with *n*, the upper limit of the CI has a peak at the point where the averaged 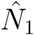 is very close to the true *N*. Near this point, the range of the CI is large and 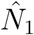 is asymmetrically distributed with a longer tail on the large side (e.g., *n* = 200 in Fig. 5c). As *n* increases beyond this point, the range of the CI decreases and the shape of the distribution asymptotically becomes to symmetric.

Next, we evaluated the accuracy of 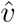. Theoretically, the bias of 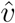 (*b*_var_) was found to have a peak at a certain value of *n*, as shown in Figs. 6a and 6b. The value of *n* at the peak decreases with the value of *c*. Figures 6c and 6d illustrate the ratio of the averaged 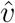 to the variance of 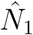, which is obtained from the IBM. If the ratio is close to unity, 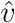 is considered an estimator of unbiasedness. There are two inconsistent results with the theoretical evaluation of 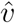. First, when *n* approaches zero, the ratio becomes inflated (e.g., *c* = 1.0 and *n* = 100 in Fig. 6c), even though *b*_var_ also approaches zero (Figs. 6a and 6b). This inconsistency may be because the values of *n* with a large ratio (*»* 1) cause the bias of 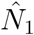. Second, as *n* increases, the ratio approaches a level less than unity, indicating a downwardly bias of the variance of 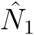. Although the degree of this bias is not very large for the investigated parameter set, this tendency is remarkable when *c* is relatively large (e.g., *c* = 3.5 and *n* = 500 in Fig. 6c). This inconsistency might result from the assumption that the correlation between pairs can be ignored and thus that the number of HSP pairs in the sample follows a binomial distribution (Equation 6).

**Figure 6.**
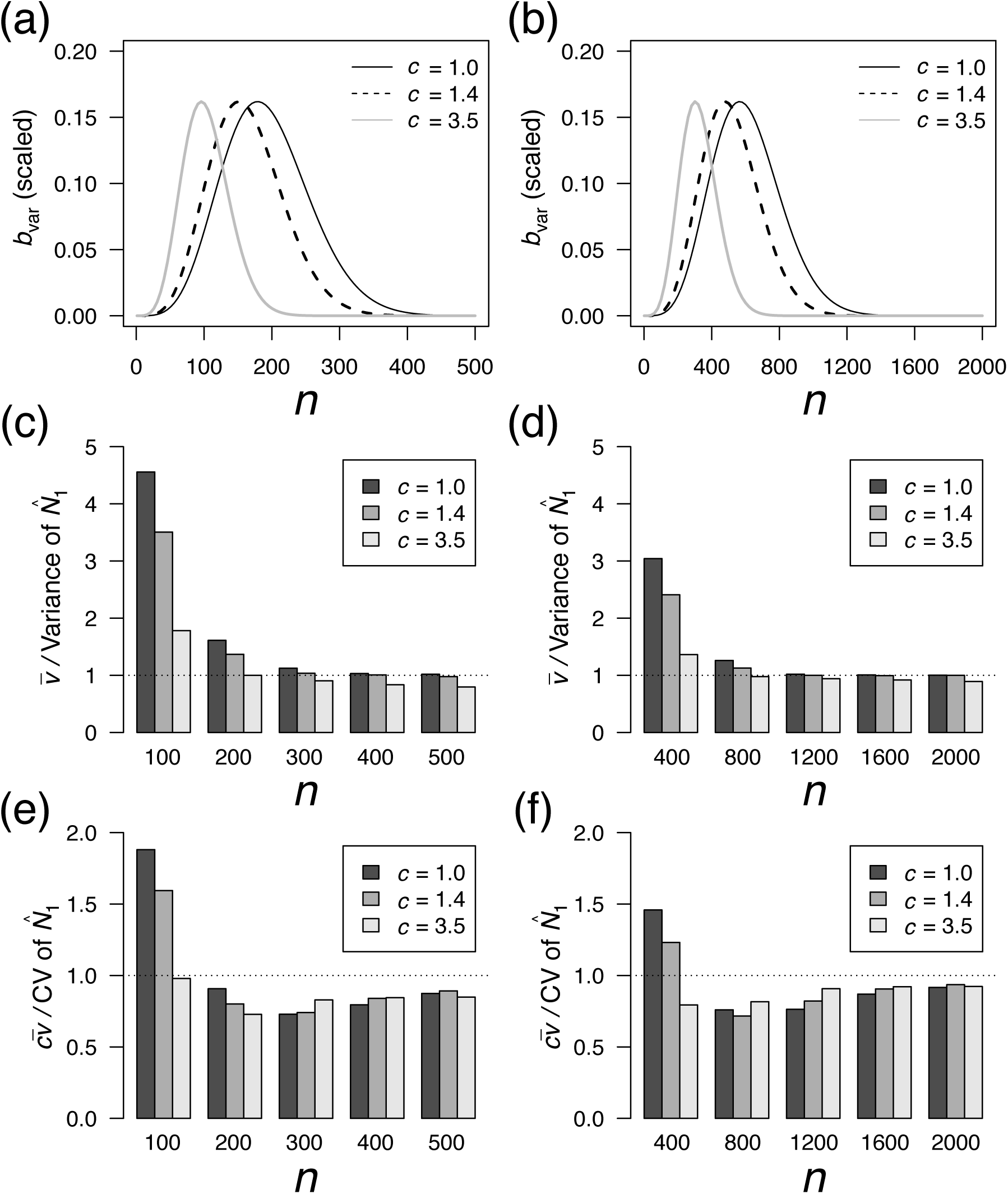
(a, b) Bias of 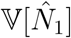 (i.e., *b*_var_) as a function of *n*. The value of the bias is scaled by (*N* + *c -* 1)^2^. (c, d) Ratio of the averaged 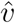 to the variance of 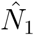 as a function of *n*. (e, f) Ratio of the averaged 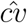 to the coefficient of variation of 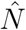 as a function of *n*. The value of *c* is described on the panel. (a, c, e) *N* = 10,000. (b, d, f) *N* = 100,000.

Finally, we evaluated the accuracy of 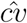 Figures 6e and 6f illustrates the ratio of the averaged 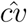to the coefficient of variation of 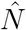, which is obtained from the IBM. Even if *n* is large enough, the ratio is less than unity, suggesting a systematic bias towards the underestimation of 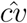 (e.g., 15% with *n* = 500 and *N* = 10,000, and 10% with *n* = 2000 and *N* = 100,000, as shown in Figs. 6e and 6f). There is no clear tendency of the degree of under-estimation among several values of *c*, implying that the cause of the underestimation is not clear.

## Discussion

We theoretically developed a nearly unbiased estimator of the number of mothers in the population 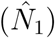, the estimator of its variance 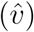, and its coefficient variation 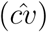, which are based on the number of MHS pairs found within the same cohort. The performance of the estimators (accuracy and precision) was quantitatively evaluated by running the IBM. Our modelling framework allowed for two types of reproductive variations, i.e., the variance of the averaged offspring number per mother (between-age variation) and the variance of the offspring number among mothers with the same reproductive potential (within-age variation); the framework summarized the two effects into one parameter (*c*). The proposed method allows us to easily extend the estimation of total adult population size when the sex ratio is given.

Our modelling framework is presented in the context of the CKMR method, which designs the kinship-oriented estimation of adult abundance. In CKMR, as far as we know, there is little theoretical foundation that can be applied to estimate the size of an adult population with overdispersed reproduction using within-cohort HSPs. The method proposed here provides information pertaining to both the number of mothers and the effect of overdispersed reproduction from the number of within-cohort MHS pairs. A previous study of the HSP-based CKMR method avoided using within-cohort HSPs, as these may lead to a biased estimation due to the effect of within-age variation (Bravington *et al.* 2016b; CCSBT 2017; Hillary *et al.* 2018). Our method overcame this limitation by directly incorporating this effect into the probability that two offspring pairs share an MHS relationship (*π*). The theoretical expectation of the number of MHS pairs (E[*H*]) derived from this probability was very accurate in matching the result of the IBM (Figs. 3a and 3b), even though the correlation between pairs was ignored (i.e., assuming a Bernoulli trial). We believe that this theoretical result provides a chance for analysing within-cohort HSPs and that it can greatly widen the scope of the CKMR method.

To estimate the number of mothers, our theoretical results provide guidance for a sample size to archive the required accuracy and precision, especially if the (order of) the number of mothers and the degree of combined overdispersion are roughly known. For example, when the number of mothers is within 10^4^–10^5^, sampling 500 offspring would fall within the range of accuracy of the estimation with a 1%–10% bias (Fig. 4). Even if there is no information about the number of mothers, the coefficient of variation of the estimated number can be estimated 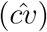when the sample size is above the appropriate level (Figs. 6e and 6f). While the estimator of the variation of the number of mothers 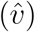 is relatively accurate for the investigated parameter set (Figs. 6c and 6d), the present estimator of the coefficient variation is systematically biased; thus, the improvement in accuracy will be an important area of future research.

Alternatively, one might expect that the number of mothers and the overdispersion parameter could be simultaneously estimated, e.g., by a maximum likelihood estimation. Equation 10 can be rewritten as

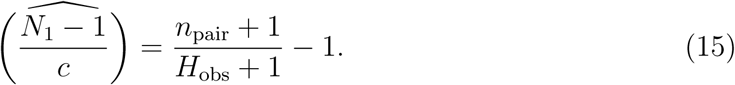

This relationship indicates that *N*_1_ and *c* cannot be estimated simultaneously from the number of observed MHS pairs. If *c* can be separately estimated, then the MHS pairs would provide a direct estimation of the number of mothers. The parameter *c* (= (1 + *ϕ*^-1^)𝔼[*λ*^2^]*/*𝔼[*λ*]^2^) indicates the degree of the combined effects of between-age variation and within-age variation. The former, which is determined by *f* (*λ*), may be influenced by population dynamics, the age of maturity, and the relationship between the mother’s weight and age (see supplementary Appendix A). Thus, we can obtain the parameter set constituting *f* (*λ*) for species for which life-history data and the relative strength of cohorts are available. The latter, which is assumed to follow a negative binomial distribution with mean *λ* and parameter *ϕ*, may be influenced by many unknown factors and is not usually directly available for estimating *ϕ*. Recently, Waples *et al.* (2018) developed a method for obtaining a minimum bound for such a parameter from fecundity data and showed that the variance is almost Poisson-like in southern bluefin tuna. However, this method does not reflect ocean stochasticity, which may cause family-correlated survivorship of offspring after spawning. Whatever the case may be, a separate estimation of *c* may not be simple for many wild species. Nevertheless, our theoretical results are useful in practical cases for the following the reason: if the effect of overdispersion *c* is invariant across years, then the left-hand side of Equation 15 may behave as an index of the number of mothers for each year. This proposed index is very informative, particularly for integrating stock assessment in fisheries management using many kinds of data (catch data and abundance index data), because it is almost independent of fishery behaviour.

On the basis of our results, we propose two approaches for estimating the degree of combined overdispersion (*c*), which is a key parameter of this work. The first approach is to estimate the number of mothers separately by other methods and to then calculate the parameter *c* from Equation 10. For example, if offspring are sampled across years and cross-cohort MHS pairs are found, the number of mothers in the more recent cohort can be calculated without the effect of within-age variation, although this method is only applied to species for which comparisons of cross-cohort pairs are possible. The second approach is to test whether the degree of overdispersion is significantly deviated from the Poisson by calculating the summary statistic, which is based on not only MHS pairs but also the mother–offspring pairs found in the sample (Akita 2018). However, this method has not yet been extended to a situation where a variation in reproductive potential exists. If the “Poisson reproduction hypothesis” is not rejected by this statistic, then *c* = 1 may justifiably be plugged into Equation 10 (note that the false-negative rate can be quantitatively evaluated); otherwise, a careful treatment of estimating the number of mothers is needed.

In this work, we separated reproductive variance into two categories: between-age variation and within-age variation, following an arbitral distribution and a negative binomial distribution, respectively. Alternatively, Niwa *et al.* (2016, 2017) theoretically showed that an offspring number distribution exhibits power-law behaviour, assuming that individual egg numbers increase exponentially and that the lifetime reproductive term distributes exponentially. In order to be compatible with our method, *π* should be replaced by *c*_*N*_, which is the coalescent probability under the power-law distribution (Niwa *et al.* 2016, 2017). It requires the estimation of the additional key parameter that determines whether the variance of the offspring number has a finite limit as *N → ∞*, likely leading to a larger uncertainty of the estimated mother size than our modelling.

Finally, we note the theoretical connection of our results to the ratio of the effective number of mothers to the number of mothers, *N*_e_*/N*. The effective number of mothers may be defined by

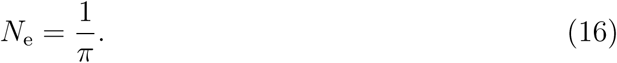

This definition is similar to the coalescent effective population size (e.g., Nordborg and Krone 2002), since the probability of sharing an MHS relationship (*π*) is identical to the probability that two individuals coalesce into a mother in the previous breeding season. Using Equations 5 and 16, the ratio can be written as

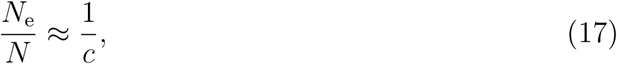

where *N » c -* 1 is assumed for the approximation. A number of literatures have shown that the estimated value of the ratio in high-fecundity marine species falls within 10^−3^– 10^−6^ (reviewed in Hauser and Carvalho 2008). Our derivation of *N*_e_*/N* based on the coalescent effective population is very simple compared to the variance effective population-based method (Felsenstein 1971), even though there is little required assumptions about the population, such as population equilibrium or stable age structure. The theoretical compatibility between our results and the existing methods for estimating *N*_e_*/N*, including methods based on linkage disequilibrium (Waples 2006), relatedness (Wang 2009), and multiple sampling (Nei and Tajima 1981), remains to be determined in future research.

## Supporting Information

Supplementary material is available at the *ICESJMS* online version of the manuscript.

## Acknowledgments

The author thanks Shota Nishijima, Hiroshi Okamura and Osamu Sakai for discussion and feedback throughout the manuscript.

## Supporting Information

### Appendix A: Probability density function and its moment of *λ*

As noted in the main text, our modelling framework does not require the specific form of *f* (*λ*) but only requires the ratio of the second moment to the squared first moment (𝔼[*λ*^2^]*/*𝔼[*λ*]^2^). However, the specific form is required for the evaluation of the theoretical results, i.e., calculating the moment or running the IBM. Here, we model an age-structured fish population, which may be a representative example showing both between-age and within-age variation.

Suppose that the mean fecundity of the mother depends on her age. Let *λ*_*a*_ be mean fecundity, which is a function of age (denoted by *a*). The moment can be described as 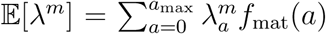 where *f*_mat_(*a*) is the frequency of mature mothers at a given age and *a*_max_ is the maximum age. Thus, we can numerically obtain the moment from *λ*_*a*_ and *f*_mat_(*a*).

For marine species with a type-III survivorship curve, it is generally assumed that individual fecundity is proportional to weight. Using the von Bertalanffy growth equation for body weight, *λ*_*a*_ is explicitly described as a function of age:

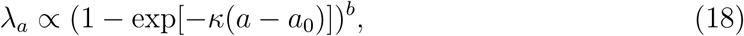

where *κ, a*_0_ and *b* are conventionally used parameters in the von Bertalanffy equation and indicate the growth rate, the adjuster of the equation for the initial size of the organism, and the allometric growth parameter, respectively. For obtaining a specific value of *λ*, a coefficient value of ten multiplied by the right-hand side of Equation 18 was used when running the IBM.

The frequency of mature mothers at a given age can be written by

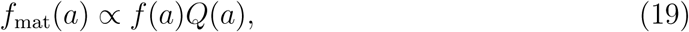

satisfying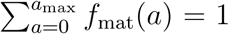, where *f* (*a*) and *Q*(*a*) indicate the frequency and maturity at age, respectively. Although *f* (*a*) is affected by historical population dynamics and age-dependent survival, for simplicity, the mortality rate is assumed to be constant (i.e., age-independent):

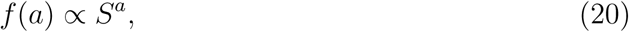

where *S* is the survival probability. Maturity at age (*Q*(*a*)) is assumed to be a knife-edge function, given by

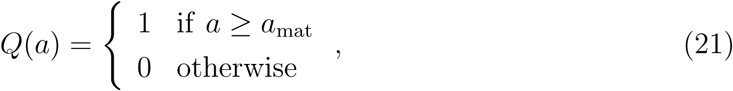

where *a*_mat_ is the mature age.

For calculating 𝔼[*λ*^2^]*/*𝔼[*λ*]^2^, the required parameter set is (*κ, a*_0_, *b, S, a*_mat_). In this paper, for the purpose of representation, we fixed the value of several parameters: *κ* = 0.3, *a*_0_ = 0, *S* = 0.5 and *a*_mat_ = 0. In addition, we chose the parameter value *c* (= (1 + *ϕ*^-1^)𝔼[*λ*^2^]*/*𝔼[*λ*]^2^) to be 1, 1.4 and 3.5 for comparison with the results in the main text, which are derived from the parameter set (*ϕ, b*) = (50, 0.0009), (5, 0.9) and (0.5, 0.9), respectively.

### **Appendix B: Properties of the moment estimator of** *N*

Here, we show that 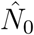 (in Equation 9) is the maximum likelihood estimator and that 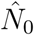 is upwardly biased, especially when *n* is small. Let *L* be the likelihood of the distribution of *H* (Equation 6). Given the observation (i.e., *H*_obs_), the partial derivative of the log-likelihood with respect to *π* is given by

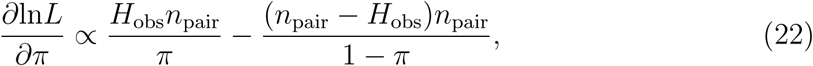

leading to the maximum likelihood estimator of *π*:

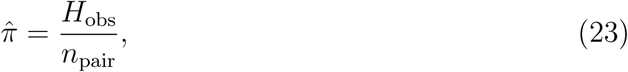

where 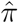 satisfies 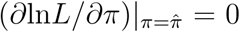= 0. Substituting Equation 5 into Equation 23, we can obtain the estimator 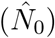 described in Equation 9.

Consider the bias of 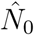 defined by 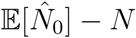. We set the following equations:

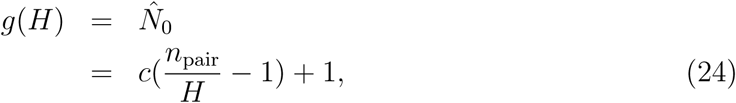

and;

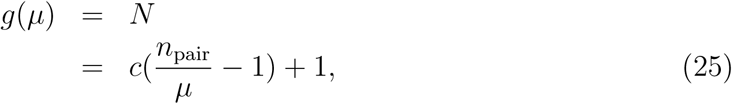

where *µ* = 𝔼[*H*]. Using Equations 7 and 8, the bias is given by

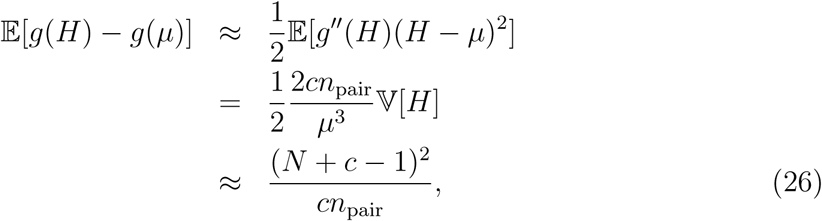

where a quadratic approximation for *g* centred at *µ* and V[*H*] *ϕ* E[*H*] are used. This result implies that *n ∼ O*(*N* ^1^*/*^2^) is required for the informative estimation from 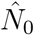 (a similar discussion appears in Bravington *et al.* (2016b)).

### **Appendix C: Derivation of the nearly unbiased estimator of** *N*

In considering,

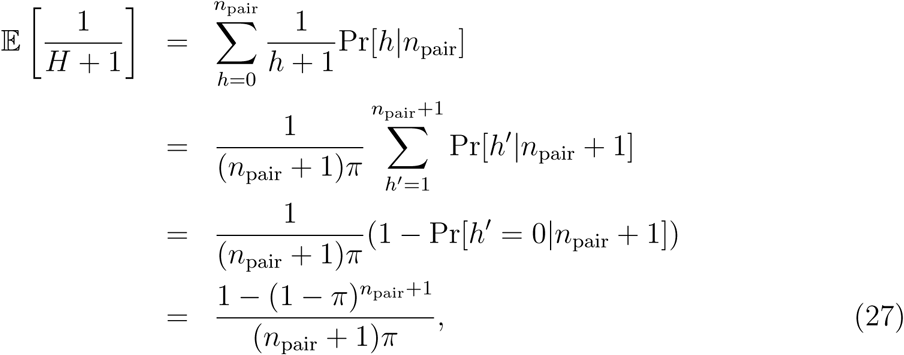

assuming the binomial form of *H* (Equation 6). Equation 27 is not directly applied for the derivation of the estimator of *N* due to the complex formulation. Thus, we simplified the formulation:

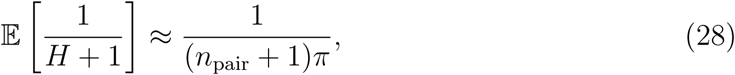

assuming

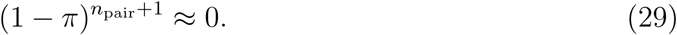

This simplification deviates from the prediction by Equation 27 when *n* is relatively small. Replacing 𝔼[1*/*(*H* +1)] by 1*/*(*H*_obs_ +1) in the left-hand side in Equation 28, we can obtain the estimator 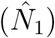 described in Equation 10.

For the evaluation of 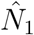, the bias is calculated. 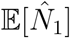 is required for the calculation and given by

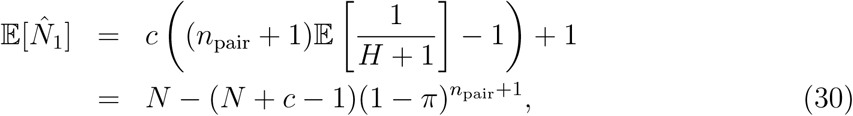

where the relationship in Equation 27 is used. This brings the formulation of the bias, as described in Equation 11.

### **Appendix D: Derivation of the estimator of the variance of** 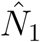

Let *v* be the estimator of variance of 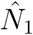. It is desirable that *v* should be defined such that the bias (denoted by *b*_var_) is reasonably small. From Equation 10, the variance of 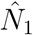 is given by

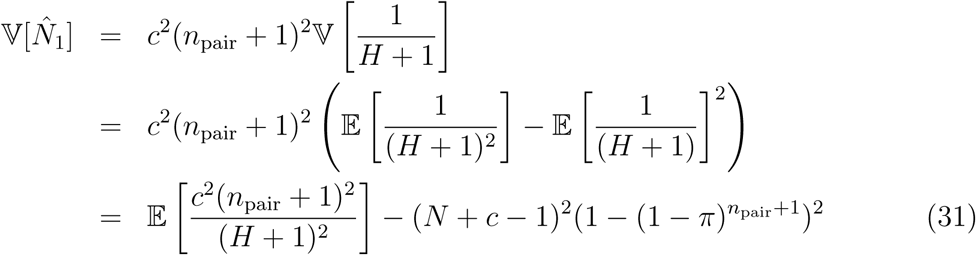

where the term E[1*/*(*H* + 1)] is calculated from the relationship in Equation 27. Roughly speaking, 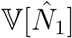 is dominated by two terms when *n*_pair_ is relatively large: 𝔼[*c*^2^(*n*_pair_ + 1)^2^*/*(*H* + 1)^2^] and -(*N* + *c -* 1)^2^. Thus, it is expected that E[*v*] includes both terms for a reasonably small bias. We proposed the following formulation for *v*:

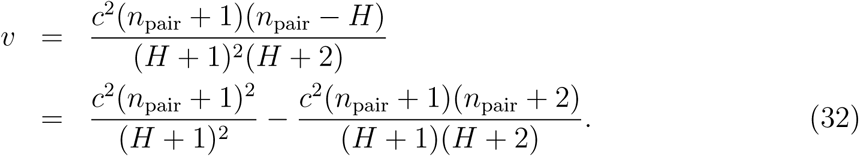

The expectation of the second term in Equation 32 is given by

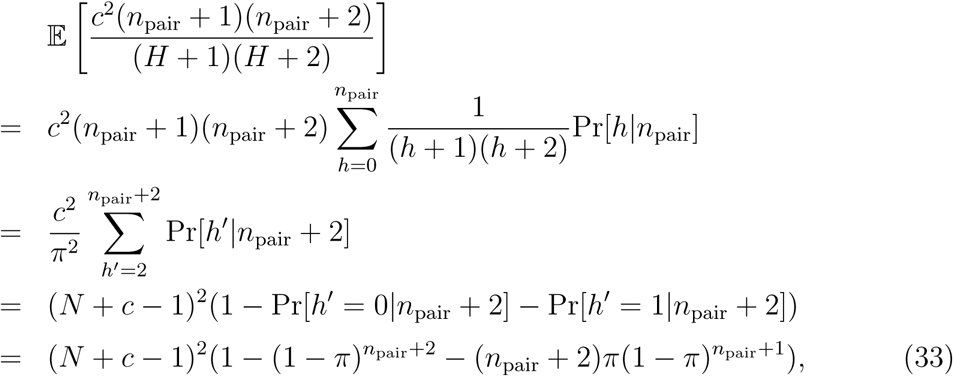

leading to a relatively small *b*_var_ when *n*_pair_ is large, which is described in Equation 13.

## References

Akita, T. 2018. Statistical test for detecting overdispersion in offspring number based on kinship information. Popul Ecol, 60.

Bravington, M. V., Grewe, P. M., and Davies, C. R. 2016a. Absolute abundance of southern bluefin tuna estimated by close-kin mark-recapture. Nat Commun, 7: 13162.

Bravington, M. V., Skaug, H. J., Anderson, E. C., et al. 2016b. Close-kin mark-recapture. Stat Sci, 31: 259–274.

CCSBT 2017. Report of the twenty second meeting of the scientific committee. 2 September 2017. Yogyakarta, Indonesia.

Chapman, D. G. 1951. Some properties of hypergeometric distribution with application to zoological census. University of California Public Statistics, 1: 131–160.

Eldon, B., Riquet, F., Yearsley, J., Jollivet, D., and Broquet, T. 2016. Current hypotheses to explain genetic chaos under the sea. Curr Zool, 62: 551–566.

Felsenstein, J. 1971. Inbreeding and variance effective numbers in populations with over-lapping generations. Genetics, 68: 581–597.

Hauser, L. and Carvalho, G. R. 2008. Paradigm shifts in marine fisheries genetics: ugly hypotheses slain by beautiful facts. Fish and Fisheries, 9: 333–362.

Hedgecock, D. and Pudovkin, A. I. 2011. Sweepstakes reproductive success in highly fecund marine fish and shellfish: a review and commentary. B Mar Sci, 87: 971–1002.

Hillary, R. M., Bravington, M. V., Patterson, T. A., Grewe, P., Bradford, R., Feutry, P., Gunasekera, R., et al. 2018. Genetic relatedness reveals total population size of white sharks in eastern australia and new zealand. Scientific Reports, 8: 2661.

Luikart, G., Ryman, N., Tallmon, D. A., Schwartz, M. K., and Allendorf, F. W. 2010. Estimation of census and effective population sizes: the increasing usefulness of DNA-based approaches. Conserv Genet, 11: 355–373.

Nei, M. and Tajima, F. 1981. Genetic drift and estimation of effective population size. Genetics, 98: 625–640.

Nielsen, R., Mattila, D. K., Clapham, P. J., and Palsbøll, P. J. 2001. Statistical approaches to paternity analysis in natural populations and applications to the north atlantic humpback whale. Genetics, 157: 1673–1682.

Niwa, H. S., Nashida, K., and Yanagimoto, T. 2016. Reproductive skew in Japanese sardine inferred from DNA sequences. ICES J Mar Sci, 73: 2181–2189.

Niwa, H. S., Nashida, K., and Yanagimoto, T. 2017. Allelic inflation in depleted fish populations with low recruitment. ICES Journal of Marine Science, 74: 1639–1647.

Nordborg, M. and Krone, S. M. 2002. Separation of time scales and convergence to the coalescent in structured populations. In Modern Developments in Theoretical Population Genetics: The Legacy of Gustave Malécot, pp. 194–232. Oxford University Press Oxford.

Ottmann, D., Grorud-Colvert, K., Sard, N. M., Huntington, B. E., Banks, M. A., and Sponaugle, S. 2016. Long-term aggregation of larval fish siblings during dispersal along an open coast. Proceedings of the National Academy of Sciences, 113: 14067–14072.

Pearse, D., Eckerman, C., Janzen, F., and Avise, J. 2001. A genetic analogue of ‘mark–recapture’ methods for estimating population size: an approach based on molecular parentage assessments. Molecular ecology, 10: 2711–2718.

Seber, G. A. F. 1970. The effects of trap response on tag recapture estimates. Biometrics, 26: 13–22.

Skaug, H. J. 2001. Allele-sharing methods for estimation of population size. Biometrics, 57: 750–756.

Skaug, H. J. 2017. The parent–offspring probability when sampling age-structured populations. Theor popul biol, 118: 20–26.

Wang, J. 2009. A new method for estimating effective population sizes from a single sample of multilocus genotypes. Molecular Ecology, 18: 2148–2164.

Waples, R. S. 2006. A bias correction for estimates of effective population size based on linkage disequilibrium at unlinked gene loci. Conservation Genetics, 7: 167.

Waples, R. S., Grewe, P. M., Bravington, M. W., Hillary, R., and Feutry, P. 2018. Robust estimates of a high Ne/N ratio in a top marine predator, southern bluefin tuna. Science advances, 4: eaar7759.

Wright, S. 1931. Evolution in mendelian populations. Genetics, 16: 97–159.

